# Hepatitis B Virus Inhibits Neutrophil Extracellular Traps Release by Modulating Reactive Oxygen Species Production and Autophagy

**DOI:** 10.1101/334227

**Authors:** Shengnan Hu, Xiaowen Liu, Ying Gao, Rongfang Zhou, Muyun Wei, Huili Yan, Yueran Zhao

## Abstract

Neutrophils, an important component of the innate immune system, release extracellular traps (NETs) to eliminate invaded pathogens by trapping and killing microbes. A dysfunctional innate immune response is a major cause of persistent hepatitis B virus (HBV) infection. HBV has been shown to reduce neutrophil responses. The objectives of the present study were to determine whether HBV influenced NETs release and to identify the underlying mechanisms. Primary neutrophils and circulating blood samples were collected from 40 patients with a chronic hepatitis B infection (CHB) and 40 healthy controls to detect NETs release using a Quant-iT Pico Green dsDNA assay and to determine the levels of HBV-DNA and HBV markers. NETs release was decreased in patients with a CHB infection, and hepatitis B surface antigen, hepatitis B e antigen and hepatitis B core antibody levels negatively correlated with NETs release. The Quant-iT Pico Green dsDNA assay and western blotting were used to examine the effect of HBV proteins (HBV X protein, HBV C protein, HBV E protein and HBV S protein) on NETs release *in vitro*. Based on the flow cytometry and western blot data, HBV C protein and HBV E protein inhibited NETs release by decreasing reactive oxygen species (ROS) production and autophagy. Overall, HBV may inhibit NETs release by modulating ROS production and autophagy to escape the immune system and promote the establishment of a chronic infection.

## Introduction

Hepatitis B virus (HBV) infection is a serious liver disease. Owing to its chronicity and the increased risk of hepatocellular carcinoma, it has become a global burden. HBV has infected 2 billion people, 360 million chronically, and leads to approximately 600,000 deaths each year(1). A dysfunctional innate immune response is a major cause of persistent HBV infection(2). Neutrophils [polymorphonuclear neutrophil granulocytes (PMNs)] are effector cells involved in innate antimicrobial defence and are also related to liver disease, since have key functions in the pathogenesis of alcoholic hepatitis(3). Neutrophils employ three major strategies to combat microbes: phagocytosis, degranulation, and the release of neutrophil extracellular traps (NETs), a process referred to as NETsosis(4), which differs from apoptosis and necrosis(5). NETs are extracellular fibrous structures composed of chromatin, histones, and several proteins, such as neutrophil elastase (NE), myeloperoxidase (MPO), cathepsin G, and proteinase 3 (PR3)(6-9). These large extracellular structures trap and kill a variety of microbes by exposing them to high concentrations of NETs-associated microbicidal factors and providing a physical barrier to prevent microbial dissemination(6, 10). In addition to their roles in infection, NETs were recently shown to have roles in various sterile diseases, such autoinflammatory and autoimmune diseases(11).

The intracellular signalling pathways that regulate NETs formation are still largely unknown. This process depends on reactive oxygen species (ROS), such as superoxide, which are generated by the NADPH oxidase Nox2(5). In addition, enzymatic activity of several enzymes such as PAD4, MPO, and NE, have been implicated in NET formation; NE and MPO synergize to drive massive chromatin decondensation before plasma membrane rupture(12, 13). According to recent studies, ROS-dependent activation of ERK and p38 MAPK mediate PMA-induced NETs release from human neutrophils(14, 15). Autophagy, essential cellular mechanism for cell homeostasis and survival, plays an important role in immunity and inflammation via pathogen clearance mechanisms mediated by immune cells, including macrophages and neutrophils(16), and has recently been reported to be required for NETs release(17, 18). Wortmannin-mediated inhibition of the autophagy activity of neutrophils leads to abnormal chromatin decondensation and prevents NETs release(19). Furthermore, the inhibition of mammalian target of rapamycin (mTOR) by rapamycin accelerates NETs release, which paralleled the increased autophagic influx(20).

HBV may reduce neutrophil responses. For example, neutrophils in patients with hepatic diseases present defects in the production of oxygen radicals(21), which are indispensable for the proper release of NETs. A significant decrease in NETs release was observed in patients with liver cirrhosis (LC), and patients with LC presenting deficiencies in NETs release display an increased rate of complications(22). Based on our data, HBV inhibits the release of NETs, which affects the function of neutrophils and interferes with the subsequent innate and adaptive immune responses against HBV, leading to the establishment of a chronic infection. Moreover, the HBV C protein (HBc protein) and HBV E protein (HBe protein) enhance mTOR activity to reduce the autophagy activity of neutrophils and suppress the ROS-dependent activation of ERK and p38 MAPK to inhibit NETs release.

## Materials and Methods

### Cases

The present study was conducted on 40 patients with chronic hepatitis B (CHB) infections (age 33.8 ±4.3 years) who met the diagnostic criteria for HBV (WS299-2008) and were recruited from Shandong Provincial Hospital and 40 healthy controls without HBV infection or autoimmune diseases (age 31.4 ±4.2 years). In addition, none of the 40 patients had been treated with antiviral drugs. All participants in this study provided written informed consent. The study protocol was approved by the ethics committee of Institutional Review Board of Shandong Provincial Hospital Affiliated to Shandong University, Jinan, China.

### Isolation of primary human neutrophils

Neutrophils were isolated from peripheral blood of donors by density centrifugation using Polymorph Prep, according to the manufacturer’s recommendations (Axis-Shield)(23). The purity of granulocytes was >97%, as determined by flow cytometry using CD15-FITC (BD Biosciences). The viability of the cells was >95%, as determined by trypan blue exclusion (Sigma-Aldrich). The obtained cells were resuspended in serum-free M-199 media.

### Quantification of cell-free DNA

Purified human neutrophils (from patients with a CHB infection and healthy controls, 5×10^5^ cells/ml) were plated on poly-L-lysine-coated wells of a 48-well tissue culture plate and incubated for 30 min at 37°C in a 5% CO_2_ atmosphere. Subsequently, cells were stimulated with 1 μM formyl-Met-Leu-Phe (fMLP, Sigma-Aldrich) for 3 h. Then, 500 mU/ml micrococcal nuclease (MNase, Thermo Fisher Scientific) were added to digest the NETs. The reaction was terminated by the addition of 5 mM EDTA (Sigma-Aldrich), and the supernatants were collected and stored at 4°C until further use. The Quant-iT Pico Green dsDNA assay was used to quantify circulating free (cf)-DNA/NETs levels, according to the manufacturer’s instructions (Invitrogen)(5, 24). The amount of DNA was reflected by fluorescence intensity and was measured at excitation and emission wavelengths of 485 nm and 530 nm, in a microplate reader (Molecular Devices Spectra Max, M2, USA).

### Serological assay for patients with a CHB infection

Serum was separated from blood samples obtained from patients with a CHB infection for HBV-DNA load tests. Levels of the HBV serum markers hepatitis B surface antigen (HBsAg), hepatitis B surface antibody (HBsAb), hepatitis B E antigen (HBeAg), hepatitis B E antibody (HBeAb) and hepatitis B core antibody (HBcAb) were quantified using the Abbot HBV quantitative test kit (Chemiluminescence Microparticle Immuno Assay) with an Architech i4000 particle chemiluminescence detection instrument. HBV-DNA levels were measured using an HBV nucleic acid quantitative detection kit (quantitative PCR-fluorescent probing method) and an Applied Biosystems 7500 fluorescence quantitative PCR instrument.

### Quantitation of NETs formation: levels of histone H3 and NE in the supernatant

Purified human neutrophils (from healthy people, 2×10^6^ cells/ml) were incubated with the HBV X protein (HBx protein), HBV C protein (HBc protein), HBV E protein (HBe protein) or HBV S protein (HBs protein) (Abcam) at 37°C in a 5% CO_2_ atmosphere, followed by fMLP (1 μM) stimulation for 3 h. Then, cells were incubated with fresh media containing DNase (40 U/ml) for 15 min at room temperature to degrade and release the NETs. The supernatant was gently removed and centrifuged at 420 g for 5 min. The cell-free supernatant was then mixed with 4× loading buffer at a 3:1 ratio before western blot analysis(25).

### Intracellular ROS assay

The intracellular ROS production in neutrophils was measured by flow cytometry using CM-H2DCFDA (Invitrogen). Neutrophils (2×10^6^ cells/ml) were incubated with HBV proteins and stimulated with fMLP (1 μM). Then, CM-H2DCFDA (5 μM) was added and incubated for an additional 30 min at 37°C in the dark. The mean cellular fluorescence intensity was quantified by fluorescence-activated cell sorting analysis.

### Autophagy detection

Autophagy activity was detected by examining the levels of the LC3 and SQSTM1/P62 proteins using western blot analysis. Neutrophils (2×10^6^ cells/ml) were incubated with HBV proteins and then stimulated with fMLP (1 μM). Subsequently, we collected cells and extracted proteins to perform western blots.

### Inhibitor studies

For analyses of signalling pathways, neutrophils were preincubated with inhibitors or DMSO for 30 min at 37°C. Rapamycin (mTOR inhibitor, CST) was freshly prepared for each experiment.

### Western blot analysis

Proteins were extracted with RIPA Lysis and Extraction Buffer (Thermo Fisher Scientific). Western blot analyses were conducted using antibodies against human histone H3 (citrulline R2+R8+R17), NE (Abcam), LC3A/B, SQSTM1/P62, phospho-p44/42MAPK (ERK1/2, Thr202/Tyr204), p44/42 MAPK (ERK1/2), phospho-p38 MAPK (Thr180/Tyr182), p38 MAPK, phospho-p70S6K (Thr389), p70S6K, phospho-mTOR (Ser2448), mTOR, phospho-ULKI (Ser757), and ULK1 (Cell Signaling Technology), followed by HRP-conjugated goat anti-rabbit secondary antibodies (Santa Cruz Biotechnology). The grey values of the strips were analysed using ImageJ software. GAPDH was used as an endogenous control for normalisation.

### Statistical analysis

The data presented here were collected from a minimum of three independent experiments with neutrophils isolated from different blood donors. The results are presented as means ±S.D. Statistical analyses were performed with SPSS software version 23. Student’s t test was used to compare differences between two groups, and correlations between the quantitative data from two groups were analysed with Pearson’s correlation tests. A P-value ≤0.05 was considered statistically significant.

## Results

### Decreased cf-DNA/NETs release in patients with a CHB infection

We assessed the isolated neutrophils of 40 patients with a CHB infection and 40 healthy controls to determine whether NETs release was suppressed in patients with a CHB infection. We used both the original values for cf-DNA/NETs and the cf-DNA/NETs-dsDNA ratio (fMLP-stimulated to unstimulated cells) to express the NETs release ability. As shown in **Fig. 1A**, the average dsDNA level in patients with a CHB infection (1552 ±160.9) was less than the level in healthy controls (2152 ±239.4), and these results were similar to the findings for the NETs-dsDNA ratio (P<0.05) (**Fig. 1B**). Based on these data, NETs release activity is suppressed in patients with a CHB infection, and HBV may inhibit NETs release.

**Figure 1.**
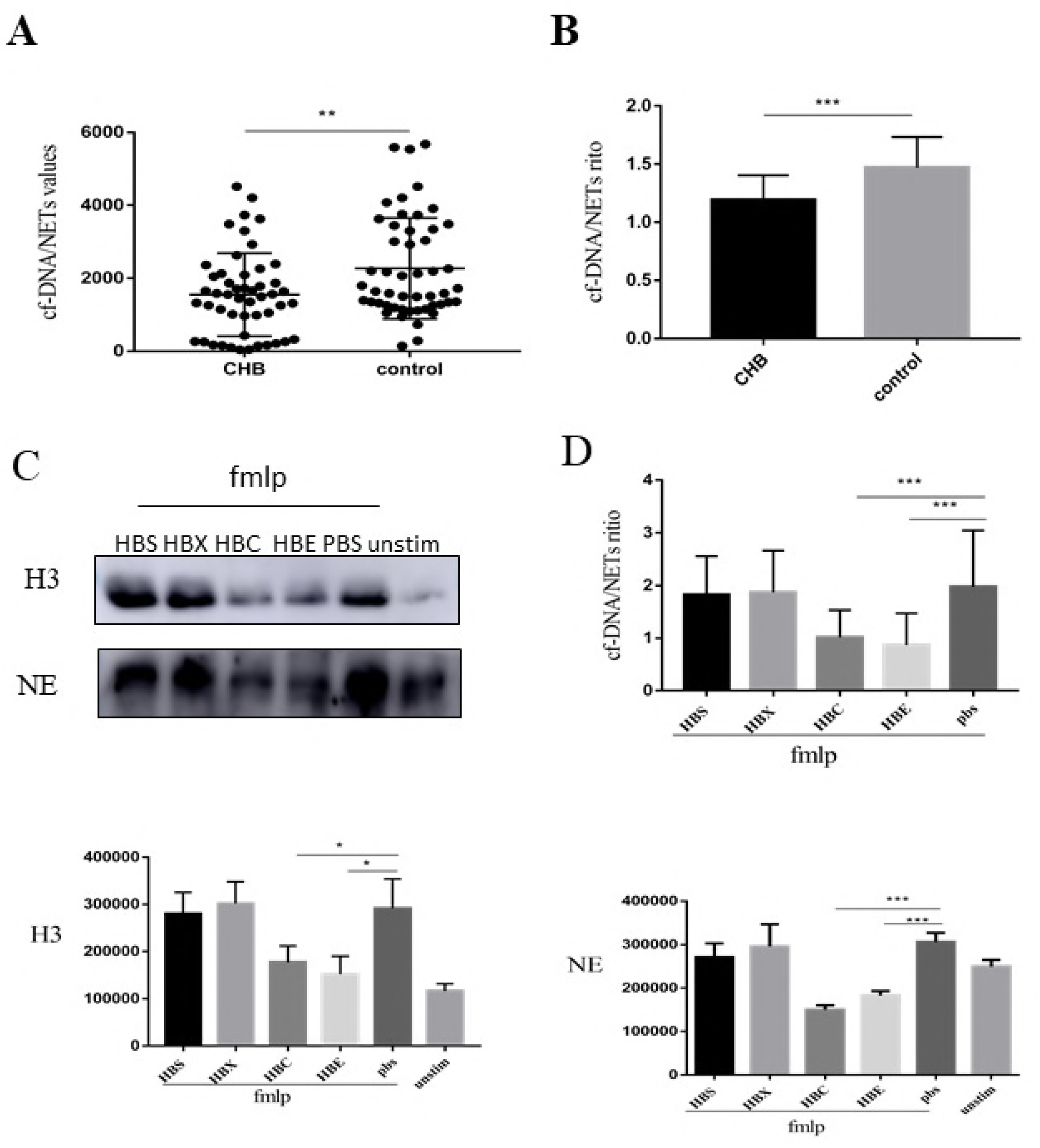
Decreased cf-DNA/NETs release in patients with a CHB infection. cf-DNA/NETs levels were quantified in the supernatants of freshly isolated PMNs from patients with CHB and healthy controls and stimulated with 1 μM fMLP for three hours. (A) The original values of cf-DNA/NETs in patients with CHB and healthy controls are shown (1552 ±160.9, n=40; 2273 ±195.5, n=40, respectively, **p<0.01). (B) The cf-DNA/NETs ratios in fMLP-stimulated and unstimulated samples from patients with CFB and healthy controls. (1.194 ±0.04113, n=50; 1.471 ±0.06153, n=40, respectively, ***p<0.001). Healthy neutrophils were treated with HBs, HBx, HBe, HBc (1 μg/ml), or PBS for 1 hour, followed by stimulation with 1 μM fMLP. (C) The cell-free supernatants were collected to quantify H3 and NE levels by western blotting (*p<0.05 and ***p<0.001). (D) After stimulation and digestion with 500 mU/ml micrococcal nuclease, the reaction was stopped with 5 mM EDTA and cf-DNA/NETs were quantified in the supernatants (***p<0.001).

### Correlations between serum levels of HBV markers and NETs

Serum levels of HBV antigens have been used in the clinic as an index of viral replication, infectivity, disease severity, and the treatment response. We determined the serum levels of HBV markers and HBV-DNA load in patients with a CHB infection to further elucidate the relationship between HBV and NETs. Then, we used a correlation analysis to determine the relationships between serum levels of HBV markers and NETs release activity. As shown in **Table 1**, the NETs ratio negatively correlated with HBsAg, HBeAg, and HBcAb levels, but no obvious correlation was observed with HBeAb and HbsAb levels. Moreover, the analysis of the correlations between the NETs ratio and HBV-DNA load did not reveal a significant correlation (**Table 2**). Thus, HBV proteins exert a complicated effect on NETs release, and different proteins have different functions.

**Table 1.**
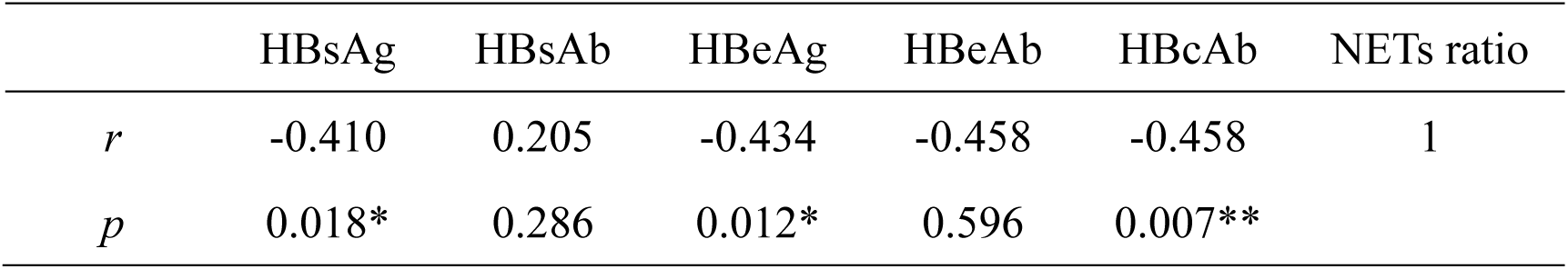
The relationship between NET release and HBV markers: (*p<0.05, **p<0.01)

**Table 2.**
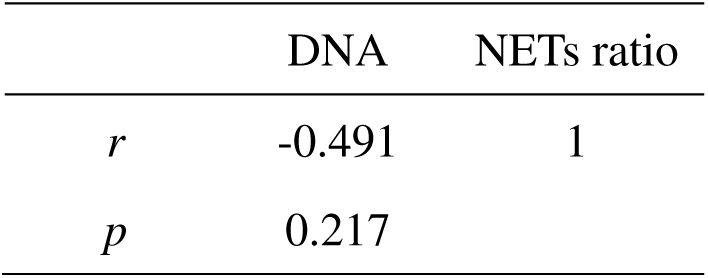
The relationship between NET release and HBV-DNA load

### The HBc and HBe proteins inhibit NETs release by suppressing ROS production and autophagy

We cultured neutrophils from healthy people with HBs, HBx, HBc, and HBe proteins and PBS to further characterise the effects of HBV proteins on NETs release. After cells were stimulated with fMLP, we measured the levels of the following markers of NETs: cf-dsDNA, histone-H3 and NE. As shown in **Fig. 1C**, in the presence of the HBc protein and HBe protein, the levels of cf-DNA/NETs were decreased, suggesting that HBc and HBe proteins inhibit NETs release, while the HBs and HBx proteins had no effect. The western blots for H3 and NE also confirmed these findings **(Fig. 1D)**.

We treated neutrophils from healthy people with CM-H2DCFDA after co-culture with HBV proteins and fMLP stimulation to determine if the diminished ROS production induced by HBV proteins directly inhibited NETs release. Since ROS were produced at high levels in each group, we used the median value to represent ROS production. According to the flow cytometry data, the HBc protein and HBe protein reduced ROS production in neutrophils (**Fig. 2A**), but the HBx protein and HBs protein did not exert significant effects.

**Figure 2.**
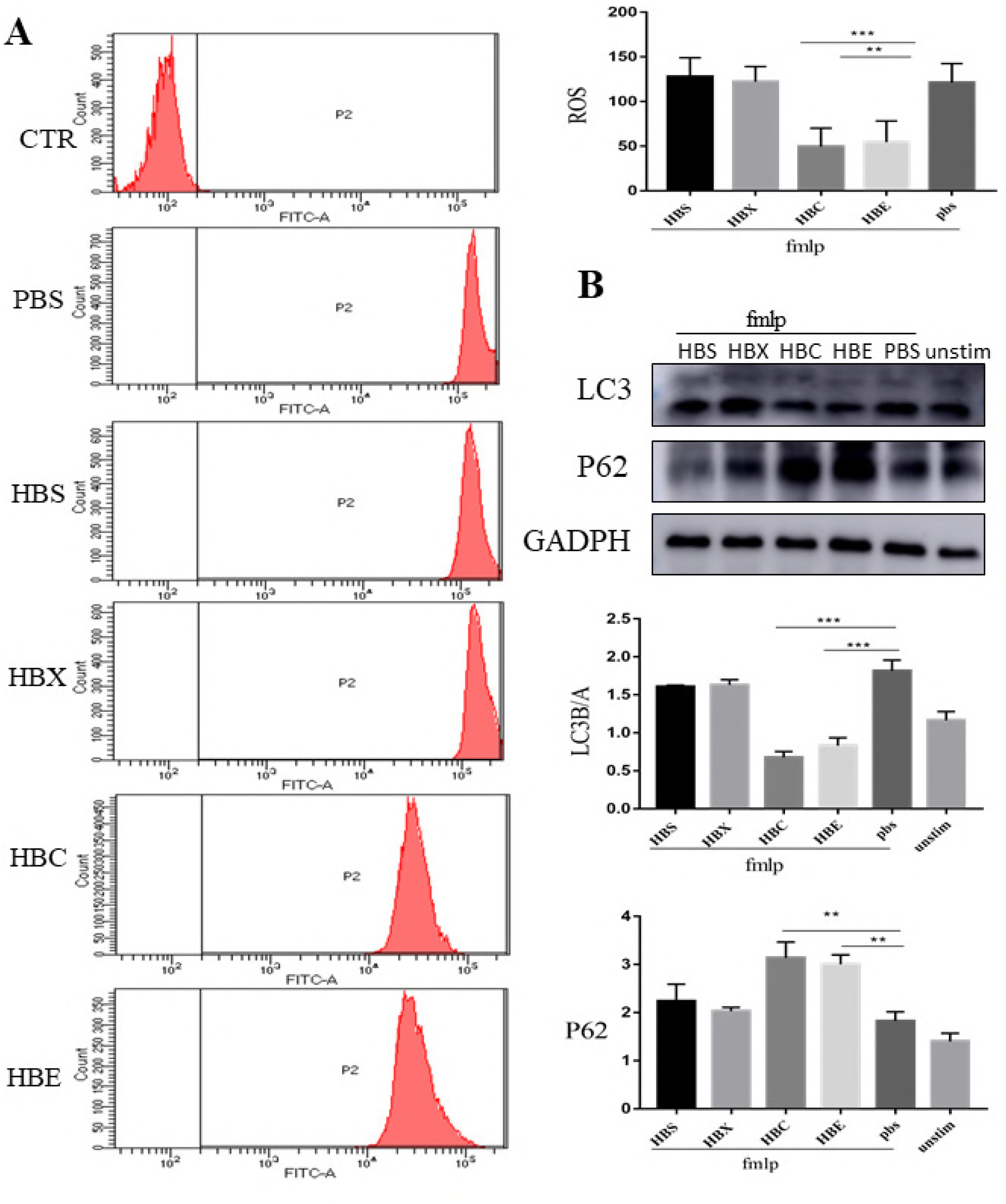
HBc and HBe proteins suppress ROS production and autophagy. **(A)** Neutrophils (5×10^5^ cells/ml) were treated with HBs, HBx, HBc, or HBe (1 μg/ml) or PBS, stimulated with 1 μM fMLP, and incubated with 5 μM CM-H2DCFDA for 30 minutes. ROS production was detected by flow cytometry (**p<0.01 and ***p<0.001). (B) Neutrophils (2×10^6^ cells/ml) were incubated with HBs, HBx, HBc, or HBe (1 μg/ml) or PBS and then stimulated with fMLP (1 μM) for 3 hours. The levels of LC3 and P62 were analysed by western blotting. Equal loading was shown by reprobing with anti-human GAPDH antibodies (**p<0.01 and ***p<0.001).

As mentioned earlier, autophagy is essential for NETs release. The level of the LC3 protein positively correlated with autophagy activity, whereas SQSTM1/P62 levels are negatively correlated. By performing a western blot analysis of the LC3 and SQSTM1/P62 proteins in neutrophils treated with HBV proteins and fMLP, we detected the impacts of HBV proteins on autophagy during NETsosis. As shown in **Fig. 2B**, the HBc and HBe proteins inhibited autophagy. Overall, the HBc and HBe proteins inhibited ROS production and autophagy, leading to a decrease in NETs release, which may be one mechanism by which HBV evades the immune system.

### Inhibition of the ERK1/2, p38 MAPK and mTOR pathways decreases NETs release

As the ERK and p38 MAPK pathways are involved in the induction of fMLP-induced ROS-dependent NETs release, phosphorylation of these molecules was assessed by western blot analysis. Significant decreases in the phosphorylation of both p38 MAPK and ERK1/2 were observed in cells treated with HBc protein or HBe protein (**Fig. 3A**). Thus, the HBc and HBe proteins inhibited the p38 MAPK and ERK1/2 pathways to reduce ROS production.

**Figure 3.**
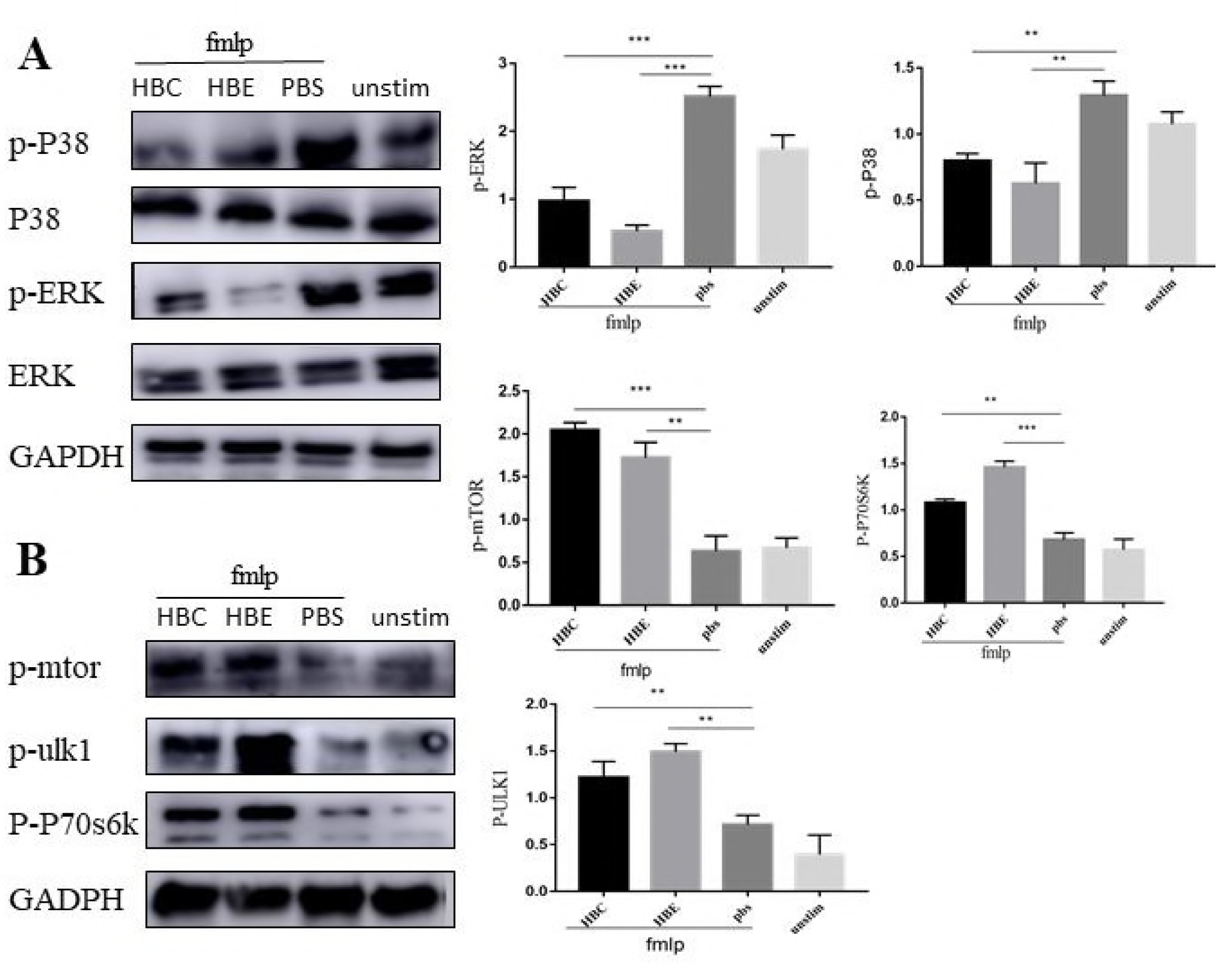
HBc and HBe protein treatments decrease the phosphorylation of ERK and p38 MAPK and mTOR pathway activation in fMLP-activated neutrophils. We extracted proteins from neutrophils treated with HBc and HBe proteins (1 μg/ml) for 1 hour and stimulated with fMLP (1 μM) for 3 hours and performed western blot analyses. (A) For quantification, the western blot signals of p-P38 MAPK and p-ERK were normalised to the GAPDH signals detected on the same blot (***p<0.001). (B) The levels of p-mTOR, p-ULK1, and p-P70S6K were detected by western blotting (**p<0.01 and ***p<0.001).

The mTOR kinase negatively regulates the translocation of proteins associated with the autophagy machinery, such as the ULK complex, to autophagy-related structures to inhibit the initiation of autophagy(18). The mTOR pathway has recently been shown to contribute to the mechanism regulating NETs formation(20). We examined the effects of the HBc and HBe proteins on the mTOR signalling pathway to investigate the mechanisms by which HBV proteins inhibit autophagy. The HBc and HBe proteins activated the mTOR signalling pathway, as evidenced by the increased levels of phosphorylated mTOR and ULK1 (**Fig. 3B**). Furthermore, we detected the levels of p-P70S6K, a protein downstream of mTOR, and observed increased levels of this protein (Fig. 4B). Based on these data, the HBc and HBe proteins may enhance the activity of the mTOR pathway and subsequently inhibit NETs release.

**Figure 4.**
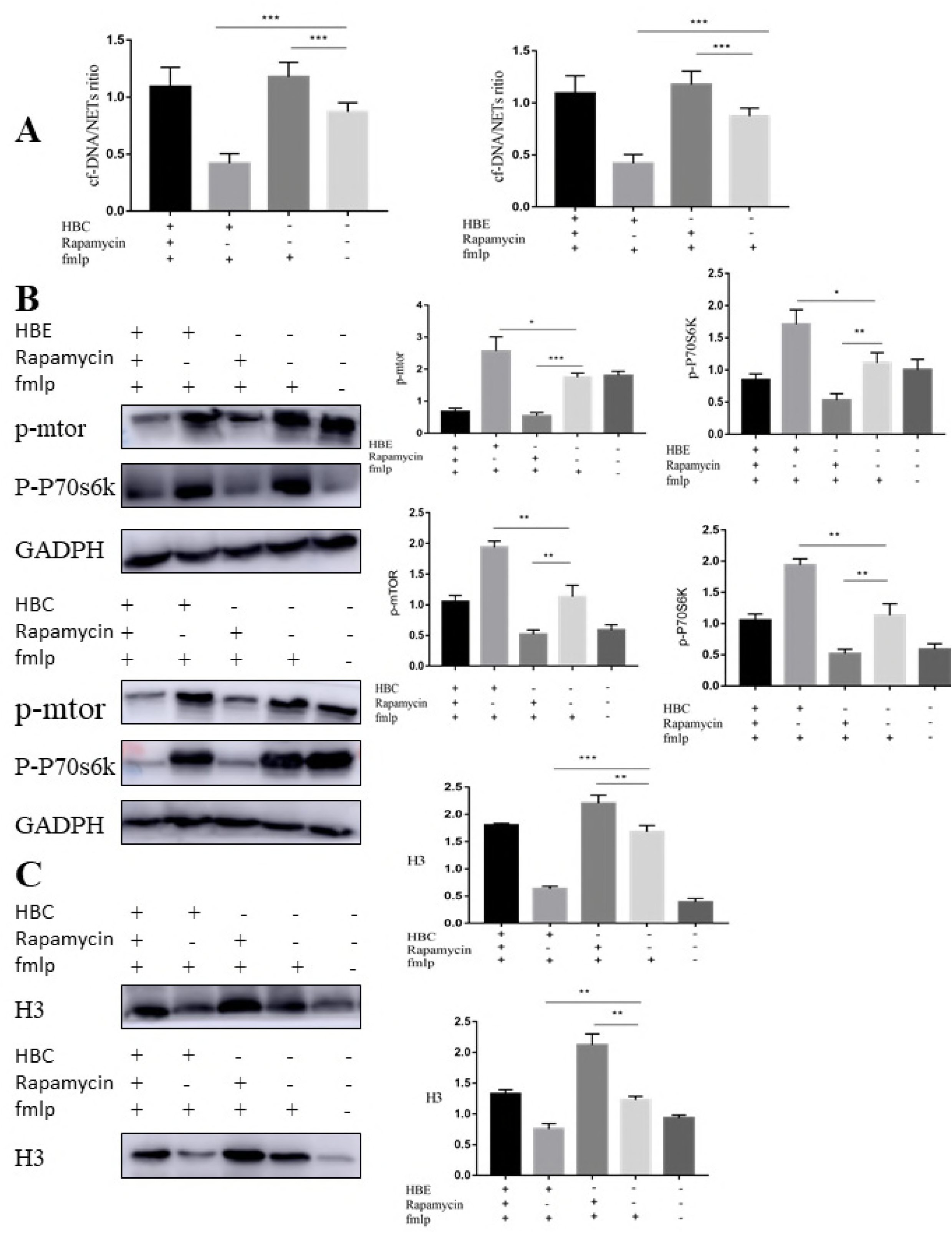
The mTOR inhibitor rapamycin increases NETs release. Neutrophils (2×10^6^ cells/ml or 5×10^5^ cells/ml) were preincubated with rapamycin (10 nM) or DMSO for 30 minutes and then incubated with the HBc or HBe protein (1 μg/ml), followed by 1 μM fMLP stimulation. (B) Levels of p-mTOR and the downstream protein p-P70S6K were detected by western blot analysis, and the results showed that rapamycin inhibited the mTOR pathway (*p<0.05, **p<0.01, and ***p<0.001). (A) cf-DNA/NETs were quantified in the supernatants. (C) The levels of H3 in supernatants were detected by western blotting (**p<0.01 and ***p<0.001).

We used the specific mTOR inhibitor rapamycin in PMNs cultured in the presence of HBc or HBe proteins to confirm the role of the mTOR pathway in the inhibitor effects of the HBc and HBe proteins on NETs release. As expected, rapamycin decreased p-mTOR levels, whereas total mTOR levels remained the same (**Fig. 4B**). Treatment with rapamycin increased H3 expression and cf-DNA/NETs levels (**Fig. 4A, Fig. 4C**), suggesting that decreased activation of the mTOR pathway is required for NETs release.

## Discussion

HBV infection remains a global health problem; however, there is still no effective measure for the cure of HBV infection. HBV persistence is often associated with the production of large amounts of viral proteins, such as the hepatitis B surface (HBsAg) and e (HBeAg) antigens(26). Neutrophils, as the foot soldiers of the immune system, play an important role in defending against various microbial infections via phagocytosis, degranulation, and the release of web-like structures called NETs.

Recent studies have examined the role of NETs release in the response to viral infections. Human immunodeficiency virus (HIV)-1 induces NETs release via the cell death pathway, and then, NETs can capture HIV virions and significantly decrease HIV infectivity. However, HIV also manipulates neutrophil activation to suppress NETs formation(27). Feline leukaemia virus inhibits NETs formation by inhibiting the activation of PKC to reduce ROS production(28). In the present study, we first showed that NETs release is decreased in patients with a CHB infection, indicating that HBV proteins suppress NETs formation and prevent the effective removal of the virus. This mechanism may be a cause of HBV persistence. These studies addressed the role of NETs in viral infection and indicated that the virus can coevolve and adapt to the innate immune response to avoid clearance.

The HBV genome contains four open reading frames (ORFs), and the core ORF encodes both the core protein (HBcAg) and the E protein (HBeAg), which share substantial sequence overlap(29, 30). HBeAg is not essential for replication or acute infection *in vivo*(31, 32) but instead has an immunoregulatory function in promoting viral persistence(33). Infants born to HBeAg-negative and HBsAg-positive carrier mothers are likely to develop acute hepatitis, but less frequently progress to the chronic disease(34, 35). Thus, HBeAg may play an essential role in HBV persistence, but the mechanisms by which HBeAg helps establish persistent infection are unclear. As shown in a recent study by Chen and coworkers, circulating HBeAg has a potential to deplete HBeAg- and HBcAg-specific Th1 cells(36), suggesting that HBeAg may regulate T cell-mediated immune responses. What’s more, based on studies performed in animal models, HBeAg may actively suppress the innate immune responses(37). We analysed the correlation between the antigens and NETs activity. Serum HBeAg levels negatively correlated with NETs release, and the HBe protein inhibited NETs release, which may be a mechanism by which HBV escapes the immune system and establishes a chronic infection.

The HBcAg is the major constituent of the nucleocapsid and is essential for viral replication and assembly(33, 38). HBcAg functions as both a T cell-dependent and T cell-independent antigen(39). HBcAg-specific cytotoxic T lymphocytes control HBV replication and the progression of liver damage(40). Because of the important role of T cells in the antiviral process and the specificity of HBcAg, most researches about HBcAg are related to T cells. Surprisingly, the serum HBcAb level negatively correlated with NETs release and the HBc protein inhibited NETs formation in the present study. Activated neutrophils modulate T cell proliferation(41). HBcAg may inhibit neutrophil activation, which leads to T cell dysfunction and ultimately enables HBV replication and a persistent HBV infection. Moreover, inhibited NETs do not effectively trap and kill the virus, which may be another reason for HBV persistence. HBsAg is the first indicator and an important biomarker for the diagnosis of HBV infection(42).

High circulating HBsAg levels in patients can contribute to the hampered immune response(43). According to the results of the correlation analysis, high HBsAg levels reduced NETs activity. Furthermore, HBsAg did not directly impact NETs activity. HBsAg was recently shown to directly contribute to the dysfunction of mDCs to escape the immune system(43). HBx plays an important role in HBV replication and infection(44), but few studies have examined neutrophils. In the present study, the HBx protein had no effect on NETs formation.

Both ROS and autophagy are required for NETsosis, as inhibition of either autophagy or ROS production prevents NETs formation(45). Based on our results, the HBc and HBe proteins suppress ROS production by inhibiting p38 MAPK and ERK phosphorylation. HBeAg suppresses p38 MAPK phosphorylation induced by TLR2 or TLR4 agonists in blood monocytes(46) and suppresses the respiratory burst in monocytes and neutrophils, which may be a cause of reduced ROS production; in contrast, no significant change was observed with HBs treatment(37). Furthermore, the mTOR pathway negatively regulates NETs formation by modulating autophagy in cells stimulated with fMLP. Our data provide evidence that the HBe and HBc proteins activate the mTOR signalling pathway to inhibit autophagy in neutrophils. Notably, treatment with an mTOR inhibitor (rapamycin) exerted a significant effect on NETs formation, further suggesting that the HBc and HBe proteins suppress NETs formation through an mTOR pathway-dependent mechanism. However, studies in human hepatoma cell lines, such as HepG2 or Huh7 cells, revealed that HBV enhances and uses autophagy to replicate its DNA, which is mediated by the X protein(47, 48). Although the HBx protein efficiently induces autophagosome formation, is the mechanism is independent of the mTOR signalling pathway(49). The persistent activation of autophagy in hepatocytes by HBV during chronic infection may play an important role in HBV pathogenesis. The inhibition of autophagy in neutrophils by the HBc or HBe protein or the induction of autophagy in hepatocytes by the HBx protein is essential for HBV replication and persistent infection.

Unfortunately, because of the sample size and methods, our study has their limits which weakening our conclusions. We only detected ROS levels and the activation of the mTOR pathway during HBV-inhibited NETs release. Many pathways and factors have potential relationships with HBV and NETs formation and should be studied. For example, NETs formation also depends on TLR2 and complement factor 3. The expression of TLR2 on peripheral monocytes was recently reported to be significantly reduced in patients with HBeAg-positive CHB compared with patients with HBeAg-negative CHB and controls(46).

In summary, HBV inhibits NETs release to escape being trapped and killed. Furthermore, HBV may reduce the responses of neutrophils, which play important roles in innate immunity and inflammation. These changes may be a mechanism by which HBV escapes the immune system. HBV may delay viral clearance and eventually facilitate the establishment of persistent infection in this manner.

## Conflict of interest

The author(s) declare that there are no conflicts of interest

## Acknowledgements

All authors have contributed significantly and approved the manuscript. Prof. Yueran Zhao designed and supervised the study. Shengnan Hu designed and performed most of the tests, analysed the data and wrote the manuscript. Ying Gao, Xiaowen Liu, Muyun Wei, Rongfang Zhou, and Huili Yan performed parts of the experiments

## Funding

This work was supported by the National Natural Science Foundation of China (31370897), Natural Science Foundation of Shandong Province (ZR2017PH035)

